# Direct conversion of somatic cells into ‘insulin-producing-cells’ by user-defined multiplex-epigenetic-engineering vector (MEEV-β)

**DOI:** 10.1101/2022.08.12.503816

**Authors:** Raza Ali Naqvi, Afsar R Naqvi, Medha Priyadarshini

## Abstract

We demonstrate here a single-step and user-friendly approach to generate insulin producing cells by gRNA driven specific-activation of PDX1, NKX6.1, MAFA, Insulin and Glut2 genes in somatic cells via multiplex-epigenetic-engineering-vector (MEEV-β) containing dCas9.P300^core^ developed by us. Sorted Glut2^+^ cells could secrete insulin in response to glucose challenge and showed expression of β-cell specific transcription factors: NKX2.2, and aforementioned genes. Expression of Cav1.3, GSK3β,, KJNC11, and SLC30A8 genes substantiated the functional insulin secreting machinery genes in these Glut2^+^ cells. Also, absence of ARX and GCG expression in these cells highlighted the specificity of the conversion.

---

Despite shortage of adequate donors islet allo-transplantation is a proven regimen to maintain long-term-normoglycemia in Type 1 diabetes (T1D) patients without risk of iatrogenic hypoglycemia^1–2^. Though the long-term survival of pig islets in monkey have already promised overcoming the allo-islets shortage^3–5^, the disguised threat of PERV virus poses another important threat yet to be resolved ^6^. Recently developed protocols for trans-differentiation of induced pluripotent stem cells (iPSCs) into β cells have also promised to render human beta cells. ^7,8^ Despite ingeniously designed-this protocol have following challenges to deal with: 1) conversion of somatic cells into iPSCs and then iPSCs into functional β-cells in seven stages, 2) different media components and/or growth factors required for each transition, 3) this approach is based on chemical-concentration-based gene induction could be sometime toxic to the cells in order to get the optimum expression to drive functional β-cell, and 4) long term cell culture (~4 weeks).

Contemplating it, we have hypothesized to formulate a single-shot and user friendly approach to convert somatic cells into β cell within 7 days. Recently, specific and directed epigenetic engineering has become possible using the RNA guided CRISPR-Cas9 system^9^. Recent studies suggested the repurposing of Cas9 via use of its mutant: dCas9 (that retains specific DNA binding potential) by fusing it with transcription activator/ transcriptional repressor.^10,11^ Due to having endogenous histone acetyl transferase activity, we have selected P300^core^ to activate of β cell specific master regulators over other strategies of repurposing of dCas9.^12^ Report revealing the activation of globin gene in absence of its bona fide transcription factor (GATA1) via promoter-specific-gRNA and dCas9.P300^core^ signifies that any gene can be activated via modification of the histones at specific location.^12^ Leveraging this basic information, we aimed at activating β cell specific genes in somatic cells via dCas9.P300^core^ mediated epigenetic engineering.

Studies involving development, characterization and induction of β cells (from iPSCs) strongly suggested PDX1, NKX6.1 and MAFA genes as master regulators essential for the induction of complex gene networking program that eventually results in functional β cells.^7,8,13–15^ Based on these reports, we prepared a schematic demonstrating the rationale of using these genes for epigenetic engineering of somatic cells in this context (**Supplementary Fig 1**). Under normal circumstances, these gene are expected to be under tight epigenetic control – located in hetero-chromatin regions in non-β cells.^15^ Therefore, to activate them in these cells, promoter specific TATA and BRE elements should be exposed for RNA polymerase II binding and onwards transcription. ^16^ To resolve this, we have mapped DNase I hypersensitive sites (nucleosome free regions) located at −100 bp of the transcription–start-site, preferably DNase hypersensitive region associated with H3K4Me1 cluster (to favor the intrinsic support for chromatin remodeling) selected for gRNA designing^17^ (**Supplementary Fig 2A-C**). Multiple gRNAs at these said sites were designed for each gene to evaluate their contribution towards gene activation (**Supplementary Table 1**). Each gRNA was separately cloned in pSPgRNA vector (**Fig.1A-C**) and co-transfected with pc-DNA-dCas9.P300^core^ in human somatic cells (**Supplementary Fig. 3**). After 4 days, gRNAs via dCas9.P300^core^ facilitated the activation of PDX1, NKX6.1 and MAFA genes. Based on RT-qPCR data, we have selected the best guides: gRNA2 (PDX1), gRNA3 (NKX6.1) and gRNA3 (MAFA) owing to their higher expression amongst all tested (**Fig. 1 F, G, H)**. Herein, we have hypothesized: if we could express the insulin and Glut2 gene in these cells, the opening of more heterochromatin regions via P300^core^ will make somatic cells more susceptible to conversion-might be due to positive feedback effect on their transcriptional factors. Importantly, Glut2^+^ β cell subtype in human pancreas was strongly associated with glucose stimulated insulin secretion (GSIS).^18^ Contemplating it, we have also designed gRNAs of DNase I hypersensitive sites associated with the promoters of said genes (Supplementary Table 1) and cloned them separately in the same way as we have cloned other master regulators of beta cells. Likewise we have selected their best guides and following are the best guides for these genes: gRNA4 (Insulin) and gRNA3 (Glut2). (**Figure 1D, E, I, J**).

**Figure 1:**
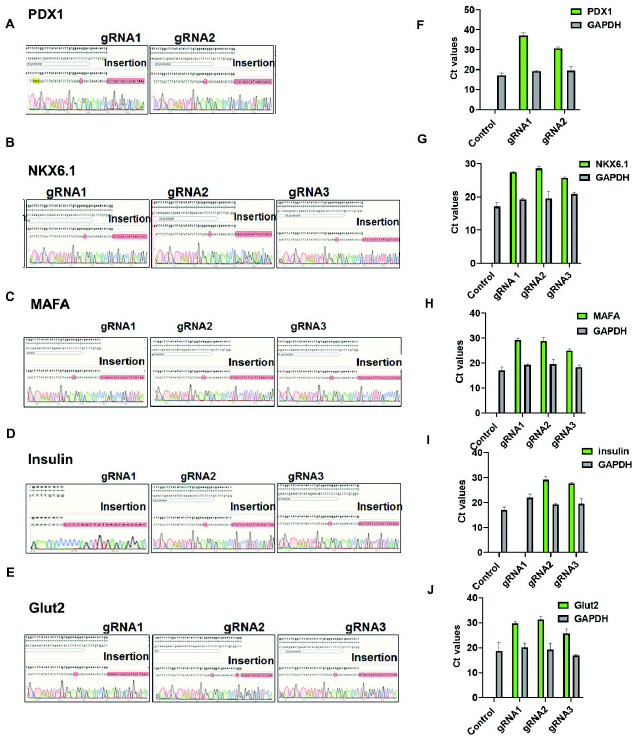
Elucidation of targeted epigenetic engineering and activation of beta cells genes in human lung fibroblasts using dCas9. P300^core^. (**A-E)** Cloning confirmation of annealed oligonucleotides targeting 5’ DNase I hypersensitive sites of PDX1, NKX6.1, MAFA, Insulin, Glut2 genes in pSPgRNA vector at BbsI sites (**F-J)** qRT-PCR of PDX1, NKX6.1, MAFA, Insulin, Glut2 driven by different gRNAs as a result of epigenetic engineering by dCas9.P300^core^..

At this point, we thought that opening the chromatin at multiple points would leave no choice or enforced somatic cells to convert into functional insulin producing cells. We had two choices in this pursuit. One strategy is to co-transfect all these five the best guides directed to the said genes (PDX1, NKX6.1, MAFA, insulin and Glut2) along with dCas9.P300^core^ vector in somatic cells. In this strategy-the likelihood of having all guides in one somatic cells are quite slim. However, second strategy is: clone all these guides and dCas9. P300^core^ in one vector and transfect it in somatic cells. Since, multiplex gene editing strategy is published for Cas9 based gene editing but it has not been done for epigenetic editing. The problem is: pX330AdCas9 1×5 vector (that has potential to possess 5 guides) is available but no dCas9. P300^core^ vector is available. Therefore, we have clone in-frame P300^core^ in pX330AdCas9 1×5 vector. The in-frame fusion of P300^core^ with dCas9 in px300A1×6 dCas9 was performed in following steps: 1) NLS (nucleus localization signal) removal from dCas9 using restriction digestion, 2) amplification of P300^core^ with NLS from pc-DNA-dCsa9.P300 vector in way its 5’ end overlaps with dCas9 and 5’ end overlaps with the 3’ end of pX330AdCas9 1×5 vector left after restriction digestion in step 1, 3) assembly of P300^core^ fragment in px330A1×5 vector (**Fig. 2A**). Correct in-frame fusion was confirmed by DNA sequencing of left and right junctions to P300^core^ and full sequence of P300^core^ in final vector. After confirming it, 1) we have cloned the best guides NKX6.1, MAFA, insulin and Glut2 in pX330S2, pX330S3, pX330S4, and gRNA for PDX1 in pX330AdCas9.P300^core^ 1×5 and 2) subjected to golden gate assembly of aforementioned vectors containing gRNAs for specific genes with pX330A dCas9.P300^core^ 1×5 (**Supplementary Fig, 4, Fig. 2 B-F**). We called this vector as multiplex epigenetic engineering vector-β (MEEV-β). Human somatic cells available in our cell bank were transfected with MEEV-β and Glut2^+^ cells were flow-sorted at day 7 (**Fig. 3A, Supplementary Fig. 5**). These cells were collected in low-binding 96 well plates and spun @ 200g for 1 min and challenged with low and high concentration of glucose consecutively to evaluate the insulin production. Sorted Glut2+ cells demonstrated insulin secretion at high (28 mM) concentration of glucose with a fold change of 1.5 (with respect to low glucose, 2.8 mM concentration) (**Fig. 3B**).

**Figure 2:**
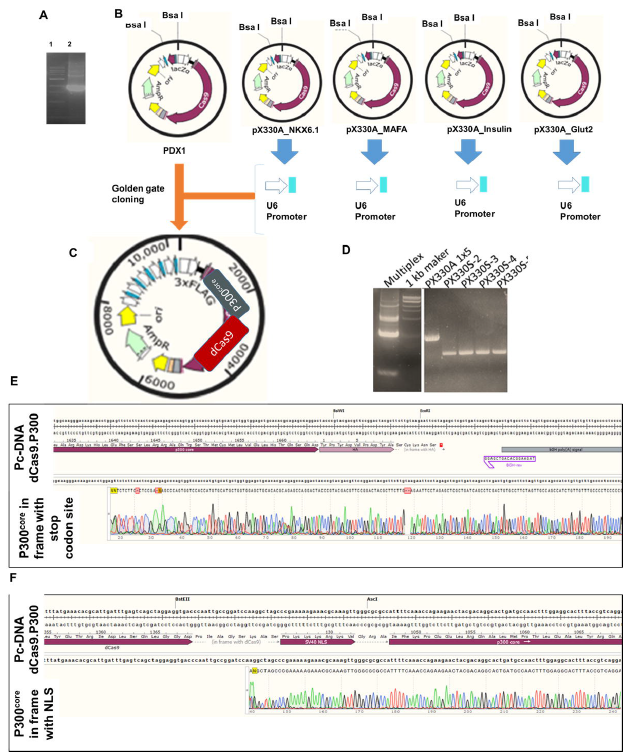
Induction of functional insulin producing cells by in-house developed multiplex epigenetic engineering vector. (**A)** Amplication of P300^core^ from pc-DNA-dCas9.P300^core^ vector containing 5’ and 3’ of double digested pX330 1×5 dCas9 vector (double digested : EcoR1 and Fse I) (**B**) Schematics of cloning of individual gRNAs targeting DNase I hypersensitive regions of PDX1, NKX6.1, MAFA, insulin and Glut2 genes in pX330 dCas9 1x 5, pX330S2, pX330S3, pX330S4 and pX330S5 vectors (**C)** Schematics of Bsa I based golden gate cloning of all gRNA targeting above regions in pX330 dCas9.P300 vector made via fragment amplified in (A) (**D)** PCR based confirmation showing presence of all five annealed oligos cloned under U6 in pX330 dCas9.P300^core^ vector. (**E and F)** Confirmation of in-frame cloning of P300^core^ fragment in addemnled vector containing all five annealed oligos by DNA sequencing. The sequence alignment was done in SnapGene. Schematics of cloning and assembly in (**B**) and (**C**) was generated in SnapGene.

**Figure 3:**
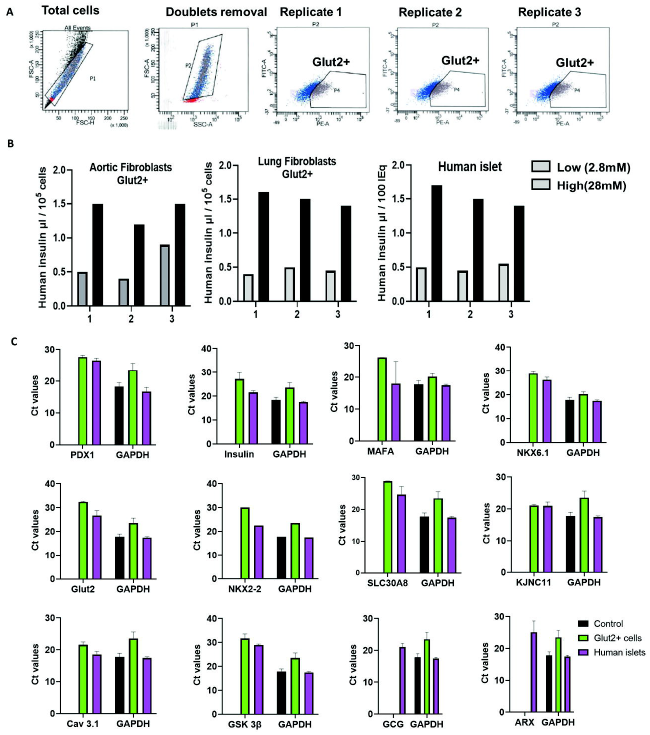
Sorting and functional analysis of Glut2+ cells. (**A**) Flow sorting showing the presence of Glut2 marker at the surface of lung fibroblasts transfected with multiplex epigenetic vector (MEEV-β). P4 gate showing the presence of Glut2+ cells. Replica 1-3 showing the sorted Glut2+ cells from three independent experiments, showing the reproducibility of the technique (**B**) Glucose stimulated insulin secretion (GSIS) by sorted Glut2+ cells collected in 96 well low binding plates. 2.8 mM was the low concentration of glucose and 28 mM was the higher concentration of glucose to ensure the dose dependent insulin secretion by Glut2+ sorted cells. 1-3 number on X-axis showing the number of times the Glut2+ cells were challenged with glucose (**C)** qRT-PCR for beta flow-sorted Glut2^+^ cells. Two replicates were used to analyze the gene expression. Each bar presented here as average ±s.d of biological replicates. Ct values on Y axis is cycle numbers after which the signals of each gene appeared during qPCR analysis. GAPDH was used as housekeeping gene. RNA isolated from 100 IEq/DNA of human islets was used as a positive control for the expression of beta cell specific genes. The expression of ARX (alpha cell specific transcription factor) and GCG (glucagon) was used as a negative control: as these are pancreatic alpha cells specific markers.

Next, we have addresses whether Glut2^+^ cells possess gene defining the β cell integrity: insulin processing and storage, specific voltage gated channel proteins needed for glucose stimulated insulin secretion. The rationale of selecting specific genes in this context are given in Supplementary Figure 6, therein we have made simple graphical representation showing the involvement of key genes in glucose stimulated insulin production storage and secretion (**Supplementary Fig. 6**) RT-qPCR data showed that Glut2+ cells not only express the targeted genes by MEEV-β but also demonstrate the expression of their down-stream targets required for β-cell development (NKX2-2), and function beta cells: SLC03A08, KJNC11, GSK, Cav1.3, (**Fig. 3D, Supplementary Fig 6,)**. RT-PCR data of Glut2+ cells was more consistent in terms of the expression of aforementioned genes, thereby recommending the use of fibroblasts for this conversion. Absence of alpha cells specific enzyme GCG (glucagon) and ARX (alpha cell specific transcription factor) in these cells suggested the specificity of our method.

Therefore, in-house generated beta cell specific all-in-one vector opened/ activated multiple beta cells specific genes native β-cells would results in activation of beta cell developmental and functional machinery (insulin production). These results ensures that direct conversion of β cells from somatic cells is very much possible. Our strategy of using multiple gene activation in a single-shot reduce the time-lag between activation of beta cell master regulators for development (PDX1, NKX6.1 and MAFA) and downstream genes (Glut2 and Insulin) in a single-shot. Importantly, the sorting of Glut2^+^ cells ensures that these cells are really glucose sensing cells that will surely make insulin upon glucose challenge. This attribute, not tested in any transplantation studies, would give confidence to the clinicians in future that they are transplanting the purified active insulin producing cells in the patients.Since only ¼th in total beta cell population in pancreas is Glut2^+18^ At this point, we also expect cell therapy using these cells might reduce the total number of beta cells required to maintain normoglycemia. All these said merits need to be tested in pre-clinical studies.

## METHODS

### Cell culture

Flow-sorted lung endothelial cells (CD31^+^), human aortic endothelial cells (CD31+) from were maintained in EGM™-2 Endothelial Cell Growth Medium-2 Bullet Kit™ (Cat # CC-3162; Lonza) and EGM™ −2 MV Microvascular Endothelial Cell Growth Medium-2 Bullet KitT^M^ (Cat #CC-3202; Lonza) in collagen coated culture dishes (Cat # 08-772-75; Corning). Lung fibroblasts were maintained in complete DMEM containing 10% FBS (Cat #95059-636; VWR), 1% Penicillin-Streptomycin (Cat #15140122, ThermoFisher Scientific) and 1% anti mycotic agent .PBMCs isolated by Ficoll® Paque Plus (Cat # 45-001-750) gradient method were maintained in complete RPMI : 10% FBS, 1% Pen-Strep, 1% anti mycotic agent. Also, PBMCs were stimulated by PHA (Cat # 00-4977-93; ThermoFisher Scientific) for 48h prior to transfection.

### Construction of targeted multiplex epigenetic engineering vector, transfection and flow sorting

All the guides for PDX1, NKX6.1, MAFA, Insulin and Glut2 (presented in **Supplementary Table 1**) was designed by online tool (http://crispr.dbcls.jp/). Having tested the best guides for the expression of each beta cell specific gene, we have cloned their respective oligonucleotides in following vectors: PDX1 in pX330A-1×5 (Plasmid #58769, Addgene), NKX6.1 in pX330S-2 (Plasmid #58778; Addgene), MAFA in pX330S-3 (Plasmid #58779, Addgene), Insulin in pX330S-4 (Plasmid #58780, Addgene) and Glut2 in pX330S-5 (Plasmid #58781, Addgene). Then, as per instructions of Multiplex CRISPR dCas9/FokI-dCas9 Accessory Pack (Addgene), we have combined them all in a pX330A-dCas9 1×6 (Plasmid #63600, Addgene). Cloning of oligonucleotides was confirmed by DNA sequencing. As per instructions of kit 1.5 ul of 100 ng/ul pX330S-2 (NKX6.1), pX330S-3 (MAFA), pX330S-4 (Insulin) and pX330S-5 (Glut2), 1.5 ul of 50 ng/ul pX330A-1×6 (PDX1), 10x T4 DNA ligase buffer (2 ul; Cat # B0202S; NEB), Bsa I (1ul; Catalog # R0535S NEB), Quick Ligase (1ul; Cat #E6047 NEB), sterile nuclease free water (8.5 ul; Ambion) were mixed for golden gate assembly reaction and incubated at following thermal cycling conditions : 37 °C for 5 min, 16 °C for 10 min for 25 cycles. After that, added Cut smart (2 ul, Cat # B7204; NEB), Bsa I (2ul) in the reaction mix and incubated at following thermal cycling conditions: 37 °C for 1h, 80 °C for 10 min, 4 °C for ∞. The vector was transformed in One Shot™ TOP10 Chemically Competent *E. coli* cells (Cat #C404010) as per manufacturer’s instructions. Purified plasmid was screen for golden gate assembly by PCR using CRISPR-step2 (**Supplementary Table 2**). For *in-frame* fusion of P300^core^ in dCas9 of pX330A dCas9 1×6 (PDX1_NKX6.1_MAFA_Insulin_Glut2) vector, NLS was removed from restriction digestion using following restriction enzymes: EcoR1 (Cat # R0101; NEB) and Fse I (Cat # R588; NEB). Simultaneously, P300core including NLS signal was amplified from pc-DNA-dCas9.P300^core^ vector using Q5 High Fidelity 2x Master mix (Cat # M0492S NEB). Primer sequences are mentioned in **supplementary Table 3**. For 50 ul reaction: 25 ul 2x Q5 master mix, 10 uM forward primer (5 uL), 10 uM reverse primer (5 uL), 200 ng pc-DNA-dCas9.P300^core^ and rest nuclease free water. Following was the temperature profile for the amplification in thermal cycler : 98°C for 30 sec (initial cycle), 98 °C for 10 sec, 72 °C for 30 sec, 72 °C for 15 min : 35 cycles, and extension 72°C for 20 min. The PCR product was eluted from gel extraction kit (Qiagen). Briefly, 50 ng linearized vector, 100 ng P300^core^ were mixed with NE Builder HiFi DNA Assembly Master Mix (Cat # E2623, NEB) and incubated at 50°C in thermal cycler for 1h. Approximately, 2.5ul assembly product was transformed in One Shot® Stbl3™ Chemically Competent *E. coli* cells (**Catalog #** C737303), to avoid the intragene homologous recombination through repetitive sequences in P300^core^ segment. To confirm correct orientation, we have designed primers for left and right junction flanking P300^core^ in assembled vector (Supplementary Table 2). After that, this vector was transfected with lung endothelial cells, lung fibroblasts, aortic endothelial cells and human PBMCs (Table 4). After 14 days, these cells were flow-sorted by anti Glut2-PE (NBP2-22218SS, Novus Biologicals).

### Quantitative RT-PCR

RNA from transfected and Glut2^+^ flow sorted cells were isolated by RNeasy Plus micro kit (Cat #74136, Qiagen). cDNA synthesis was done by iScript™ cDNA Synthesis kit (Cat #1708891, BIO-RAD) in each condition. Total 20 ul reaction volume was used for cDNA reaction. The reaction mix was kept in thermal cycler with following temperature profile: 25°C for 5 min (priming), 46 °C for 20 min (reverse transcription), 90 °C for 1 (reverse transcriptase inactivation), and hold at 4 °C. Realtime PCR was performed using Sso Advanced™ Universal SYBR, (Cat # 1725271, BIO-RAD) and a CFX96 Real-Time PCR Detection System with a C1000 Thermal Cycler (BIO-RAD). Primers for qRT-PCR are listed in Table 4.

### Glucose Stimulated Insulin Secretion

Sorted Glut2+ cells were plated in collagen coated 96 well plates (1000 cells each well). After 48h cells showed the signs of attachment and the cells were carefully washed in Krebs buffer containing 2mM glucose. Following are the components of the stock solution of Krebs buffer: 128 mM NaCl (Cat #S65886; Sigma), 5 mM KCl (Cat #6858-03; Sigma), 2.7 mM CaCl_2_ dihydrate (Cat# 223506; Sigma-Aldrich), 1.2 mM MgCl_2_ (Cat #7791-16-6; J.T. Baker), 1 mM Na_2_HPO4 (Cat # S9763), 1.2 mM KH_2_PO4 (), 5 mM NaHCO3 (Cat #S6296; Sigma), 10 mM HEPES (Cat # 15630080; Life Technologies;), and 0.1% BSA (Proliant; 68700) in deionized water. After washing, the cells were incubated for 2hr in 2 mM glucose (Cat#G7528; Sigma) prepared in Krebs buffered. The cells were then washed again and incubated sequentially under following conditions: 30 min each with 2.8 mM, 28 mM, 2 mM, 28 mM, 2 mM, and 20 mM glucose in Krebs buffer. The insulin secretion was from these supernatants was detected as per instructions of Human Ultrasensitive Insulin ELISA (ALPCO Diagnostics; 80-INSHUU-E01.1).

## Acknowledgements

The authors are thankful to Wach fund grant to RAN by University of Illinois and NIH/NIDCR R01DE027980 to AN for the funding support of this manuscript.

## Authorship contribution statement

**Raza Ali Naqvi**: Conceptualized, written and corrected the manuscript. **Medha Priyadarshini:** Corrected the final draft. **Afsar Raza Naqvi**: Conceptualized and corrected the final draft.

## Declaration of competing interest

The authors have no conflict of interest.

## Supplementary information

### Supplementary Text and Figures

Supplementary Figures 1–6

Supplementary Table 1: List of oligonucleotides DNase I hypersensitive region targeting 5’ to exon 1.

Supplementary Table 2: List of primers for qRT-PCR primers

Supplementary Table 3: List of primers for PCR

## Notes

### Competing Interest Statement

The authors have declared no competing interest.

