## Supplementary Tables and Figures for "Direct conversion of somatic cells into ‘insulin-producing-cells’ by user-defined multiplex-epigenetic-engineering vector (MEEV-β)"

**University of Illinois, USA, Chicago**

**Supplementary Table 1: List of oligonucleotides DNase I hypersensitive region targeting 5’ to exon 1**

|  | **Location** | **+/- strand** | **DNase I hypersensitive site score** | **Chromosomal location of DNase hypersensitive site** |
| --- | --- | --- | --- | --- |
| **PDX1** | | | |  |
| **gRNA 1** | -276 to – 298 bp | + | 62 | [chr13:28493746-28494295](https://genome.ucsc.edu/cgi-bin/hgTracks?hgsid=797083773_wvAooG4anasT1ygdsNK3xqLR7jtx&db=hg19&position=chr13%3A28493746-28494295) |
| **gRNA2** | -521 to – 543 bp | - | 62 |  |
| **NKX6.1** | | | |  |
| **gRNA1** | -107 to -129 bp | + | 77 | [chr4:85419146-85420215](https://genome.ucsc.edu/cgi-bin/hgTracks?hgsid=797083773_wvAooG4anasT1ygdsNK3xqLR7jtx&db=hg19&position=chr4%3A85419146-85420215) |
| **gRNA2** | -492 to - 514 bp | - | 77 |  |
| **gRNA3** | -755 to -777 bp | + | 77 |  |
| **gRNA4** | -349 to -371 bp | - | 95 | [chr4:85420226-85420775](https://genome.ucsc.edu/cgi-bin/hgTracks?hgsid=797083773_wvAooG4anasT1ygdsNK3xqLR7jtx&db=hg19&position=chr4%3A85420226-85420775) |
| **Glut2** | | | |  |
| **gRNA1** | -127 to - 149 bp | - | 92 | [chr3:170745821-170746515](https://genome.ucsc.edu/cgi-bin/hgTracks?hgsid=797083773_wvAooG4anasT1ygdsNK3xqLR7jtx&db=hg19&position=chr3%3A170745821-170746515) |
| **gRNA2** | -433 to - 455 bp |  | 92 |  |
| **gRNA3** | -432 to - 454 bp |  | 92 |  |
| **gRNA4** | -1 to - 23 bp | + | 64 | [chr3:170746526-170747055](https://genome.ucsc.edu/cgi-bin/hgTracks?hgsid=797083773_wvAooG4anasT1ygdsNK3xqLR7jtx&db=hg19&position=chr3%3A170746526-170747055) |
| **Insulin** | | | |  |
| **gRNA1** | -9 to - 31bp | + | 1 | [chr11:2184006-2184155](https://genome.ucsc.edu/cgi-bin/hgTracks?hgsid=797083773_wvAooG4anasT1ygdsNK3xqLR7jtx&db=hg19&position=chr11%3A2184006-2184155) |
| **gRNA2** | -98 to - 120 bp | - | 1 |  |
| **gRNA3** | -96 to - 118 bp | + | 1 |  |
| **gRNA4** | -93 to – 115 bp | - | 1 |  |
| **MAFA** | | | |  |
| **gRNA1** | -4 to -26 bp | - | 125 | [chr8:144512921-144513215](https://genome.ucsc.edu/cgi-bin/hgTracks?hgsid=797083773_wvAooG4anasT1ygdsNK3xqLR7jtx&db=hg19&position=chr8%3A144512921-144513215) |
| **gRNA2** | -170 to - 192 bp | + | 125 |  |
| **gRNA3** | -90 to - 112 bp | + | 57 | [chr8:144513626-144514255](https://genome.ucsc.edu/cgi-bin/hgTracks?hgsid=797083773_wvAooG4anasT1ygdsNK3xqLR7jtx&db=hg19&position=chr8%3A144513626-144514255) |
| **gRNA4** | -204 to - 226 | + | 57 |  |

*DNase I hypersensitive site score = number of cell types showed this site as a DNA hypersensitive site. 1(a) and 1(b): these two sites are only reported in pancreas.*

| **Gene** | **Forward Primer (5’ to 3’)** | **Reverse Primer (5’ to 3’)** | **Accession Number** | **Specificity** |
| --- | --- | --- | --- | --- |
| **PDX1** | CCCATGGATGAAGTCTACCAAAGC | AAGTTCAACATGACAGCCAGCT | NM_000209.4 | β-cells |
| **NKX6.1** | ACACGAGACCCACTTTTTCCGGA | CTTCTTCCTCCACTTGGTCCGG | NM_006168.2 | β-cells |
| **MAFA** | CTTCAGCAAGGAGGAGGTCATCC | TCTCCTTGTACAGGTCCCGCT | NM_201589.4 | β-cells |
| **Insulin** | AGCCTTTGTGAACCAACACCTG | CACGCTTCTGCAGGGACC | NM_001185098.1 | β-cells |
| **Glut2** | GCTACCGACAGCCTATTCTAGTGG | TCGCCCTGCCTTCTCCACA | AH002747.2 | β-cells |
| **CHGA** | CTCTGAACACAGGCAGCTTTCT | ATGTTACAGTCAGGAGTTCTCAGCTTTC | NM_001275.4 | β-cells |
| **NeuroD1** | TTGCACCAGCCCTTCCTTTGATG | TCGCTGCAGGATAGTGCATGGTAA | AK313799.1 | β-cells |
| **NKX2-2** | TGAACTCTACGCCGTGTTTACAGAATG | GACATTAACGCTGGGACGGTTT | NM_002509.4 | β-cells |
| **ABCC8** | ACCACAGCACATGGCTTCATTTC | TGTACAGGTGCAGATGGTGGGATT | NM_001287174.2 | β-cells |
| **NEUROG3** | TAAGAGCGAGTTGGCACTGAGCAA | TTTGAGTCAGCGCCCAGATGTAGT | NM_020999.4 | β-cells |
| **SLC30A8** | ACAGCCAAGTGGTTCGGAGAGAAA | TTGGGAAACTGACGGTGTGACTGA | NM_173851.3 | β-cells |
| **KCNJ11** | GCGCTTTGTGCCCATTGTA | TTGATGGTGTTGCCAAACTTG | NM_000525.3 | β-cells |
| **PCSK1** | AGCTGGACCTTCATGTGATACC | GCTAGCCTCTGGATCATAGTTGG | NM_000439.5 | β-cells |
| **GAD65** | TTCCTCCAAGCTTGCGTACT | ACCATGCGGAAGAAATTGAC | NM_000818.2 | β-cells |
| **CACNA1D** | GGTGATCCCCTTCCCCATTC | ATAGTTTGCCTCGTTCGCGT | NM_001128840.3 | β-cells |
| **Kv2.2** | AGGGCAGTGTGGGCTCTTC | ATGGTAAATGTCTTGCCTACAGTTGT | NM_004770.2 | β-cells |
| **GCK** | CTTCCCTCAGTTTTTCGGTGG | TTGATTCCAGCGAGAAAGGTG | NM_001354800.1 | β-cells |
| **GCG** | AGGCAGACCCACTCAGTGAT | TCGCCCTGCCTTCTCCACA | NM_002054.5 | α-cells |
| **GAPDH** | CACCAGGGCTGCTTTTAACTCT | GAGGGATCTCGCTCCTGGAAGA | NM_002046.5 | q-RT-PCR control |

**Supplementary** **Table 2: List of primers for qRT-PCR primers**

**Amplicons obtained from all these primers are ≤ 200 bp.*

**PDX1 (pancreatic and duodenal homebox 1), NKX6.1 (NK6 homeobox 1), MAFA (v-maf musculoaponeurotic fibrosarcoma oncogene homolog A), Glut2 (carrier family 2 : facilitated glucose transporter, member 2), CHGA (chromogranin A), NEUROD1(neuronal differentiation 1), NKX2.2 (NK2 homeobox 2), ABCC8 (ATP-binding cassette, sub-family C : CFTR/MRP), NEUROG3 (neurogenin 3), SLA30A8 (solute carrier family 30 : zinc transporter), GCG (glucagon), GAPDH (glyceraldehyde-3-phosphate dehydrogenase), KCNJ11 (Potassium Inwardly Rectifying Channel Subfamily J Member 11), PCSK1 (Proprotein Convertase Subtilisin/Kexin Type 1), GAD65 (*glutamic acid decarboxylase*), CACNA1D encodes* *Cav1.3 (voltage-gated calcium channel), Kv2.2 (Voltage-gated potassium channels), GCK (Glucokinase), MAFb (v-maf musculoaponeurotic fibrosarcoma oncogene homolog b), GCG (glucagon).*

**Supplementary Table 3**: **List of primers for PCR**

| **Name of gene** | **Forward primer (5’ to 3’)** | **Reverse Primer (5’ to 3’)** | **Purpose** |
| --- | --- | --- | --- |
| **P300 with 50p overlap** | tcaccggcctgtacgagacacggatcgacctgtctcagctgggaggcgacccaattgccggatccaaggctagcc | ctggcaactagaaggcacagtcgaggctgatcagcgagctctaggaattctcaagaagcgtagtccggaacgtcgtac | For amplification of P300 |
| **Left junction primers** | Atctgggagcccctgccg | Agctgagggtccacaggttgacg | For orientation of P300 at left side |
| **Right junction primer** | GTCGGGATGCGTTTCTCACGCTGG | ATTCTCTTCCCAATCCTCCCCCTTGCTG | For orientation of P300 at right side |
| **U6 F** | ----------------------------------------------- | CTAGAGCCATTTGTCTGCAGAATT | To confirm oligonucleotide insertion |
| **CRISPR-step2** | GCCTTTTGCTGGCCTTTTGCTC | CGGGCCATTTACCGTAAGTTATGTAACG | Multiplex assembly |

**Underlined nucleotides indicating the 50 bp overlap with pX330A 1x6 dCas9 vector*

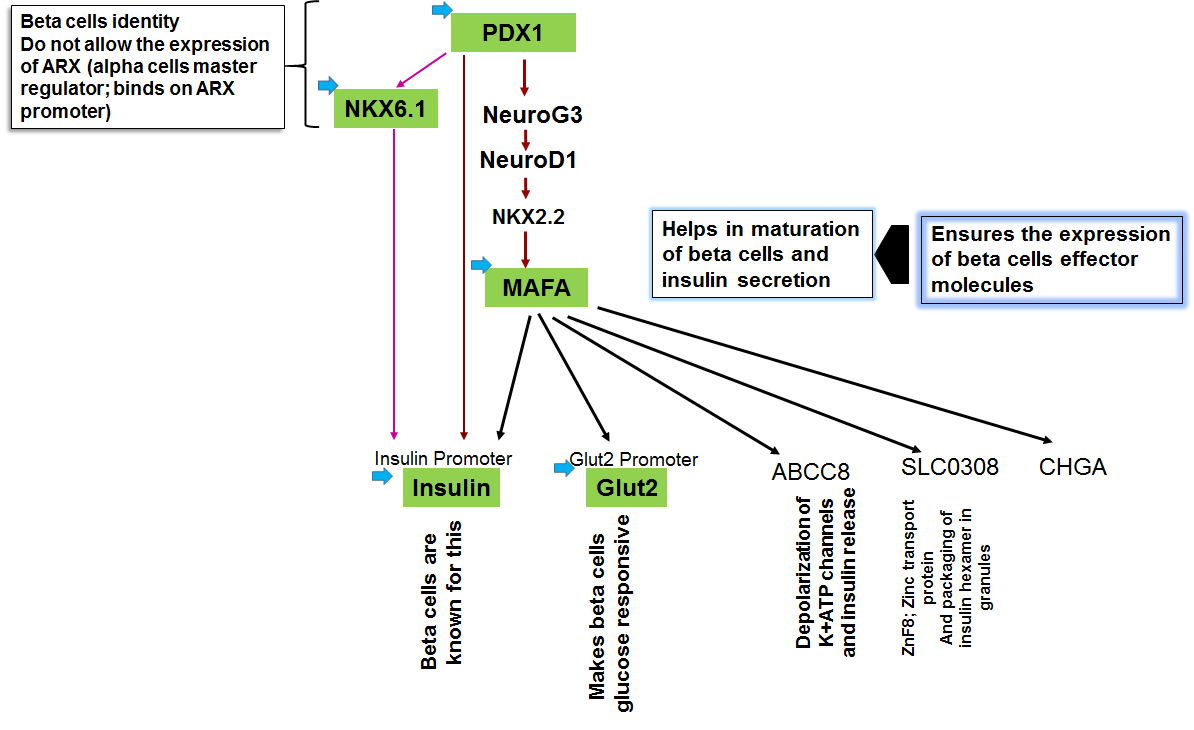

**Supplementary Fig 1. Rationale of choosing PDX1, NKX6.1, insulin, Glut2 and MAFA to induce beta cells.** PDX1 expresses in all endocrine cells initially and later on it’s only restricted in beta cells of endocrine pancreas^1-3^. PDX1 importantly induces the expression of NKX6.1- a transcription factor that restricts the development of beta cells^4^. NKX6.1 reported to bind at the promoter of ARX1 – a master regulator of alpha cells and also suppresses the glucagon expression ^5^. On other hand NKX6.1 also binds at insulin promoter and facilitates the expression of insulin^6^. Thus, NKX6.1 can function as both transcriptional activator as well as transcriptional repressor. Initially, we have thought of taking NKX6.1 to induce the beta cells genetic program, but insulin promoter- the major effector molecule of beta cells not only has NKX6.1 binding site but also PDX1 binding site^1-3^. Therefore, even though NKX6.1 can be expressed by dCas9.P300^core^ even in the absence of its transcription factor PDX1 but to get the adequate and sustained expression of insulin gene we need to have PDX1 along with NKX6.1. Also, PDX1, induces the genetic program leading to another transcription factor of insulin gene-MAFA^1^, which in turn leads Glut2 gene expression- a glucose transporter^9.10^. Learning the lessons from stem cells derived beta cells, we came to know the expression of MAFA only possible when PDX1 would induce NKX2.2 via NEUROG3 and NEUROD1^1-3^. Therefore, to reduce the time from PDX1 to MAFA, we have already targeted MAFA, so that it could provide positive feed-back for its own expression. Targeting of insulin and Glut2 simultaneously can only enforce the somatic cells to move towards the conversion of beta cells with no further decision making process related to the beta cells specific genetic network. Blue arrow indicates the activation of these loci by gRNA driven dCas9.P300^core.^

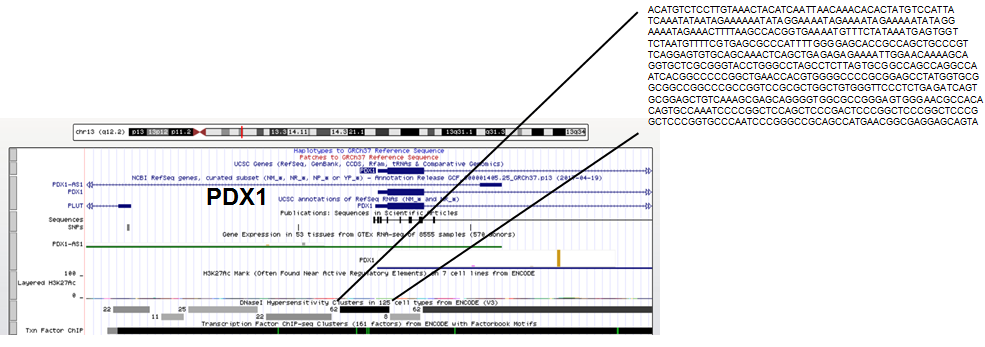
**1a**

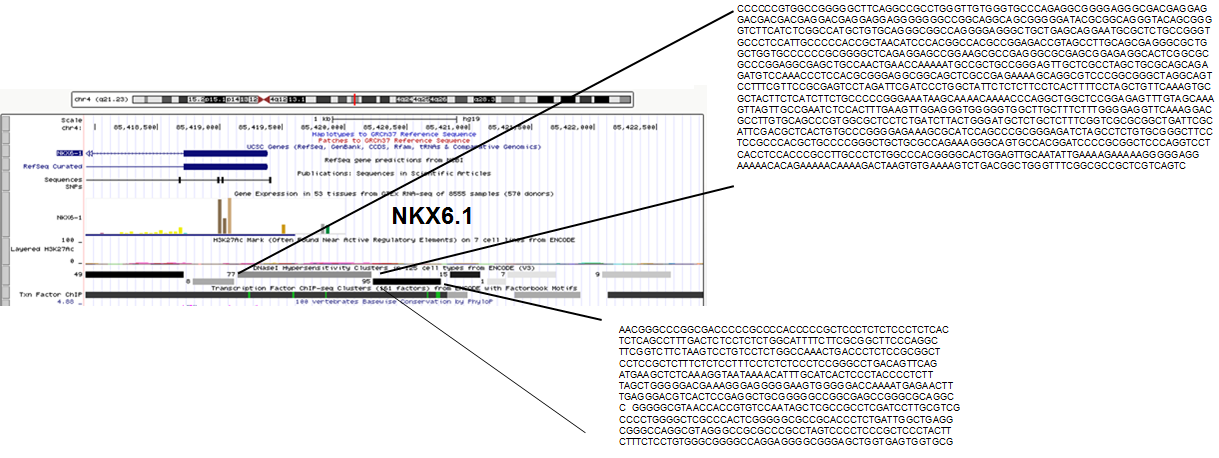
**1b**

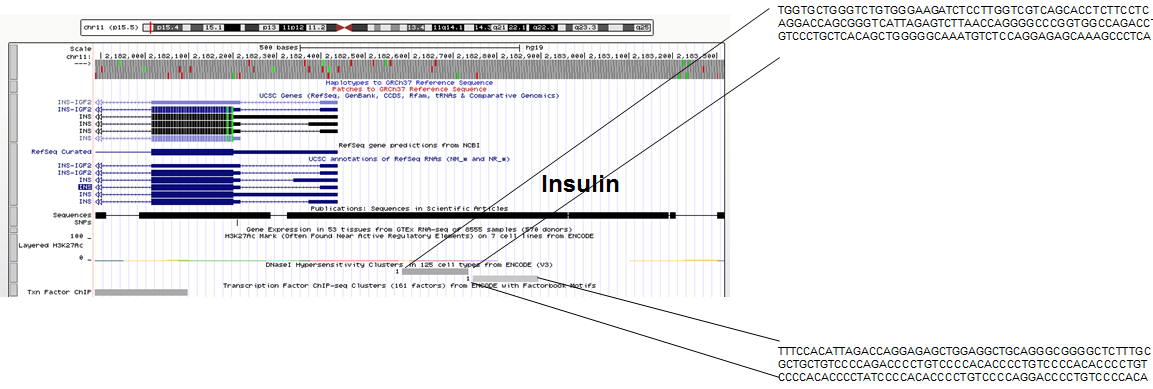
**1c**

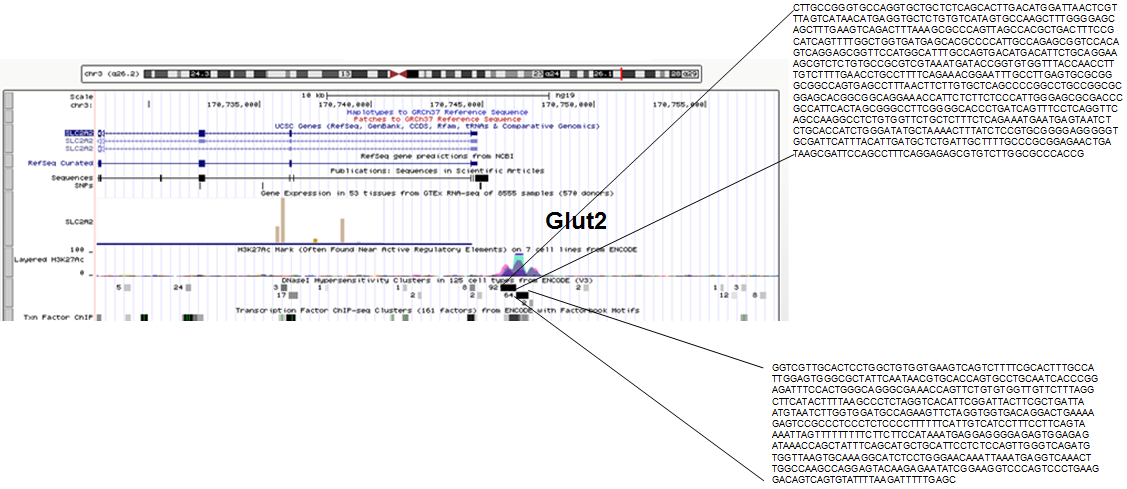
**1d**

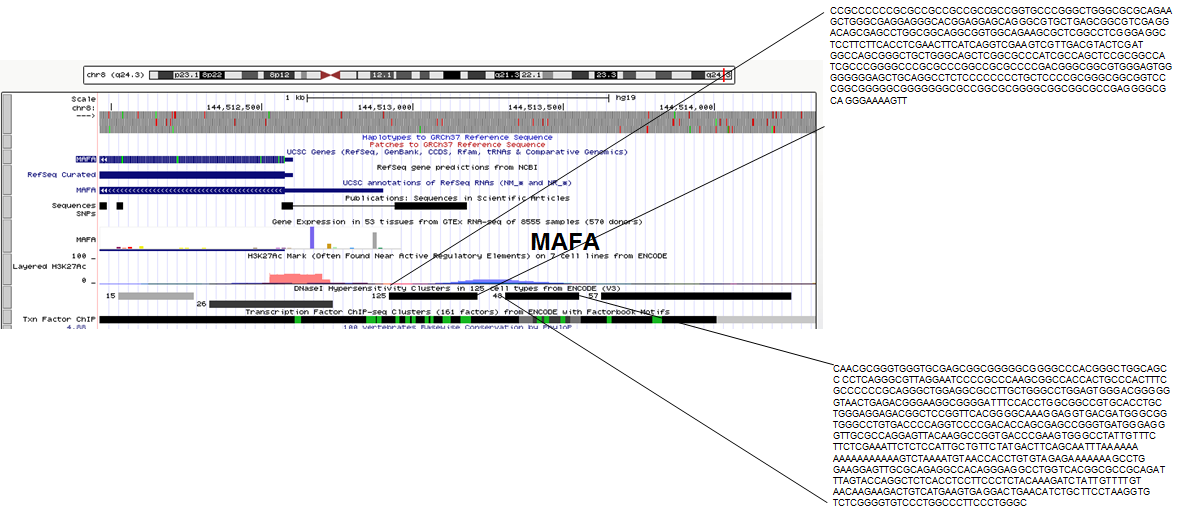
**1e**

**Supplementary Figure 2:** Mapping of DNase 1 hypersensitive region 5’ to transcription start site (TSS) of β-cells genes using UCSC genome browser. DNA sequence extraction at (a) PDX1, (b) NKX6.1, (c) Insulin, (d) Glut2, (e) MAFA genes for designing of gRNAs. Source: [**https://genome.ucsc.edu/cgi-bin/hgGateway?hgsid=782633553_HLlSFvwgjA1juVNq9lmKBHMn9gx3**](https://genome.ucsc.edu/cgi-bin/hgGateway?hgsid=782633553_HLlSFvwgjA1juVNq9lmKBHMn9gx3)**.** Number given along with DNase I hypersensitive site indicate the number of cell line showing this DNase I hypersensitive site.

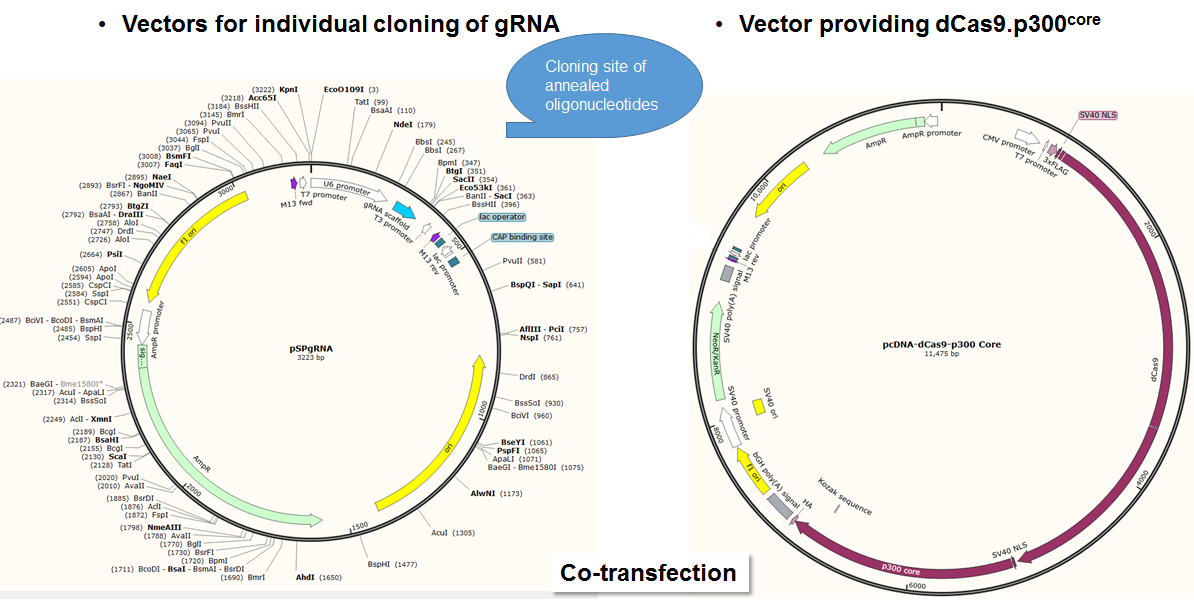
**a	b**

**Supplementary Figure 3: Snap gene generated maps of (a)** pSPgRNA vector (Addgene # 47108) with restriction sites **(b)** pcDNA-dCas9.P300^core^ **(**Addgene # 61357**).** Individual annealed oligonucleotides of each gene were cloned separately in Bbs I site of pSPgRNA vector. All the oligonucleotides (obtained from IDT) were dissolved in nuclease free water to get 100 uM concentration. ~1 ul of both 100 uM of forward and reverse oligonucleotides were mixed in 1 ul of 10x polynucleotide kinase (PNK) buffer (NEB), 1 ul PNK (NEB) and 4 ul nuclease free water. This mixture was kept in thermal cycler with following temperature cycle: 95^o^ C for 5 min, slow cooling till 25^o^ C by reducing 1 C per cycle. Cloning was done as per instructed in step1; Multiplex CRISPR/Cas9 Assembly System Kit protocol (Addgene #). Cut Smart Buffer and Bsb I was obtained from NEB. Each vector containing oligonucleotide sequence was co-transfected in lung endothelial cells along with pc-gRNA-dCads9.p300 (Addgene # 61357). These cells were in culture over a period of 14 days and then RNA was isolated by Rneasy mini kit and SYBR green based qRT-PCR was done to test the best guide

a	**b**	**c**

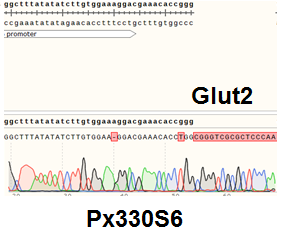

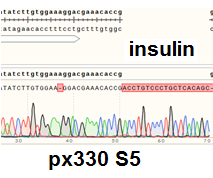

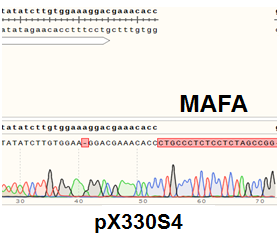

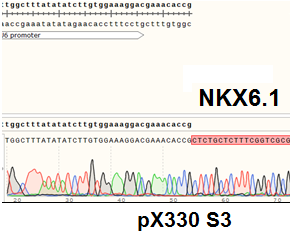

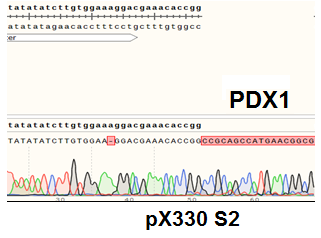
**d	e**

**Supplementary Figure 4: Cloning of annealed oligonucleotide showed comparatively better expression in donor vectors. (a)** PDX1 (gRNA2) in pX330S2**, (b)** NKX6.1 (gRNA3) in px330S3, **(c)** MAFA (gRNA3) in pX330S4, **(d)** Insulin (gRNA3) in pX330S5, **(e)** Glut2 (gRNA3) in pX330S6. All the sequencing data was analyzed in Snapgene.

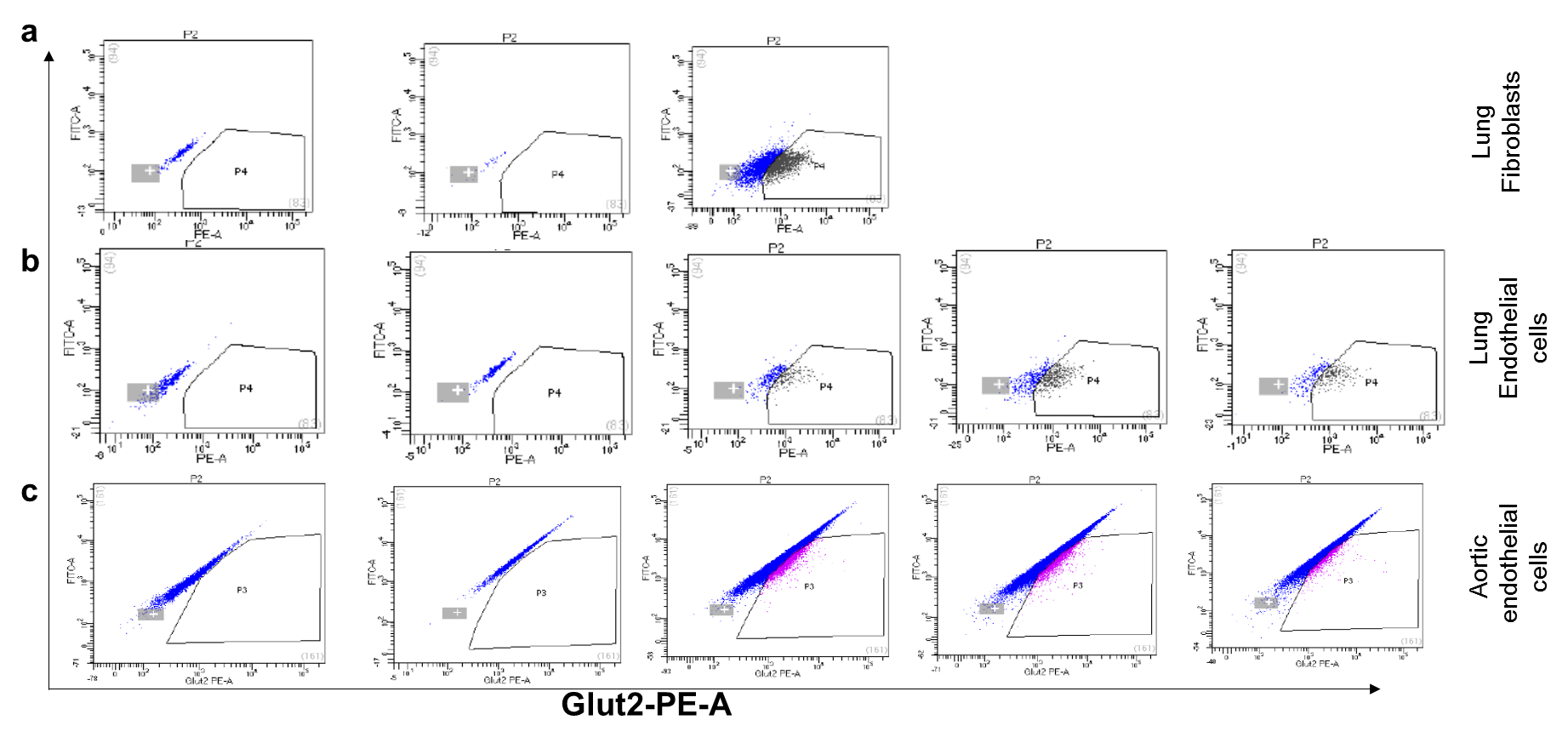

**Exp#1	Exp#2	Exp#3**

**Supplementary Figure 5: Sorting of various Glut2+ cells upon transfection with multiplex epigenetic vector in 14 days.** Three experiments were performed per cells, Exp#1, Exp#2, and Exp #3. ~1x 10^6^ cells per transfection were taken into consideration. Gate P3 in (a) and Gate P4 in (b) and (c) depicts the Glut2^+^ cells.

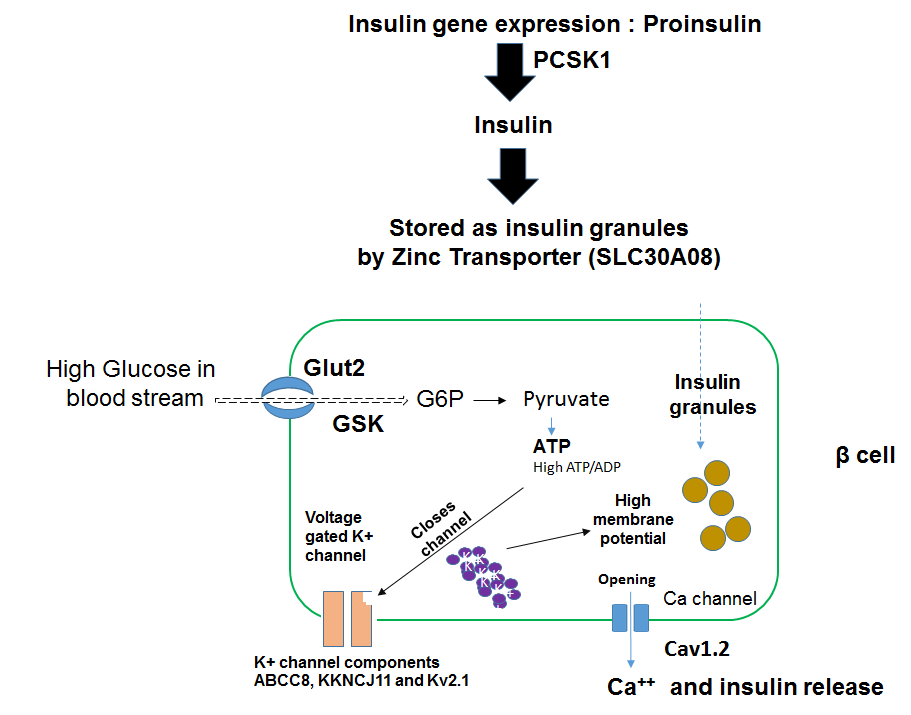

.

**Supplementary Figure 6:** Schematics of genes involved in insulin secretion. The concerted actions of PDX1, NKX6.1, MAFA, NKX2-2, NEUROG1 orchestrates the glucose responsive (Glut2+) β cells in the pancreas to produce insulin ^1-10^. Under the effect of these transcriptional factors, insulin gene transcribes and translates into pro-insulin. PCSK1 converts pro-insulin into mature insulin molecules^11^. SLC30A8 – is Zinc transporter 8 protein, mainly expressed in pancreatic islet cells and mediates zinc (Zn^2+^) uptake into secretory granules for the storage of insulin molecules^12^. The glucose independent transporter molecule SLC2A2 (also known as GLUT2) transports glucose inside beta cells^13^, which is immediately phosphorylated in glucose 6 phosphate (G6P) by glucokinase (GCK)^14,15^. This reaction is called glycolysis- results in the 1) synthesis of 2 ATP molecules, 2) and results in increased ATP/ADP ratio inside the beta cells ^13^. ATP molecule binding with ATP-sensitive K+ (K_ATP_) channels in the beta cell membrane triggers the K+ channel closure. Increase in intracellular K+ ions leads to generation of bursts of action potentials and exocytosis of insulin granules via opening of voltage gated [Ca^2+^] channels. ABCC8, KKNCJ11 and Kv2.1 are the key components of K_ATP_ channels ^14-18^.

**Supplementary Figure 9: expression of beta cell genes in transfected and sorted Aortic endothelial cells**
